## Supplementary Materials for "Defining the cell and molecular origins of the primate ovarian reserve"

**The PDF file includes:**

Figs. S1 to S8
Tables S1 to S4

**Other Supplementary Materials for this manuscript include the following:**

Data S1 to S3

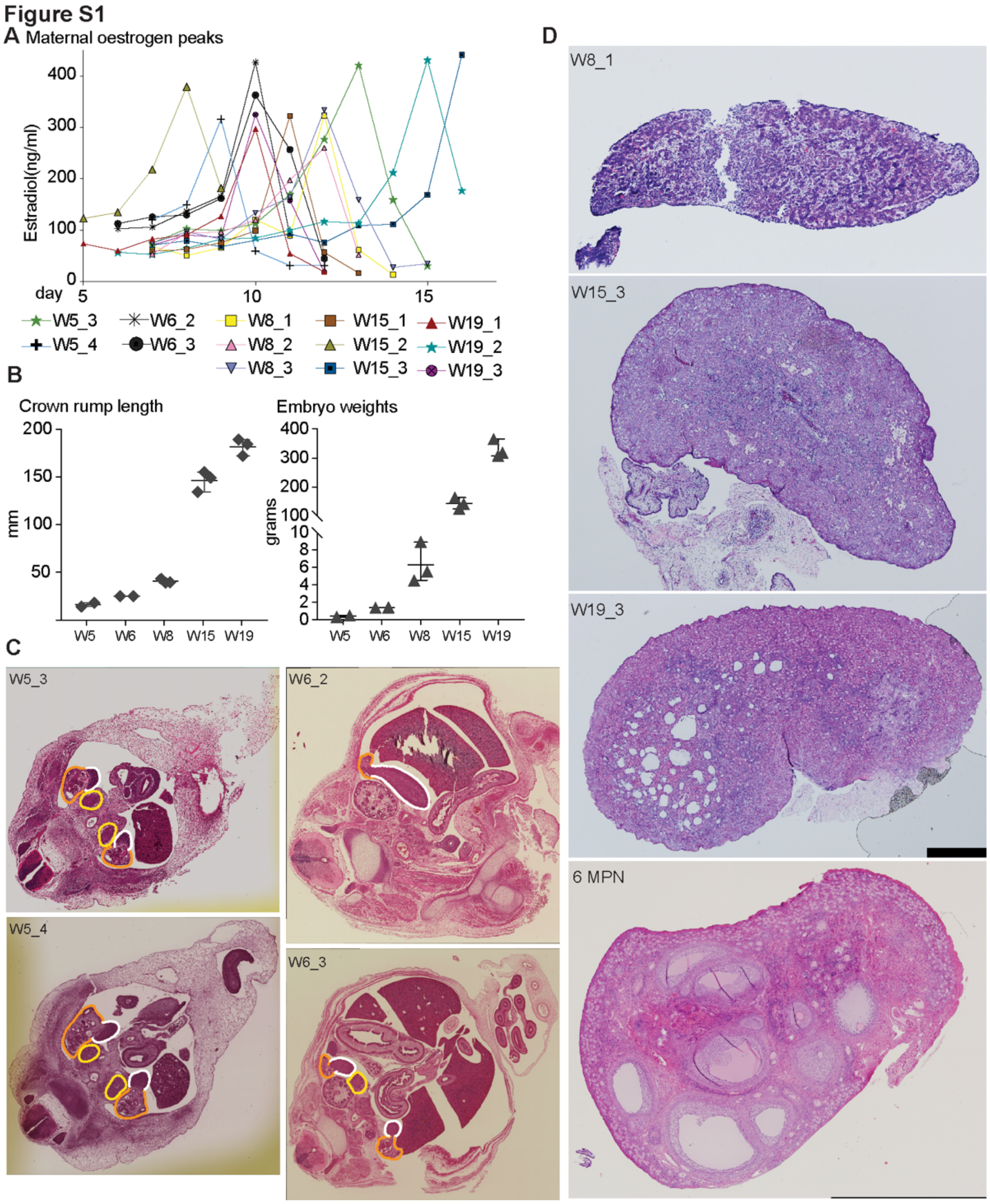

**Fig. S1. associated with Fig.1. (A)** Estradiol measurements from female rhesus macaques used to determine the timing of the estrogen peak prior to pairing with a male for time mated breeding to collect embryo samples for this dataset (n=13 animals). Day 1 (D1) of embryo development was estimated to occur 72 hours after the estradiol peak. Maternal data listed by corresponding embryo moniker. **(B)** Crown rump length and weight measurements of samples collected for this dataset (n = 13). **(C)** Hematoxylin and eosin (H&E) stain of W5 and W6 embryo torso sections. Ovary (white outline), mesonephros (orange) and adrenal (yellow, where present) regions are outlined. **(D)** H&E stain of one representative gonad dissected out at CS23, D100 and D130; replicates confirmed to be equivalent. Due to tissue size, images were compiled from multiple images using the Stitching plugin in Fiji imaging analysis software (*91*).

**Figure S2**

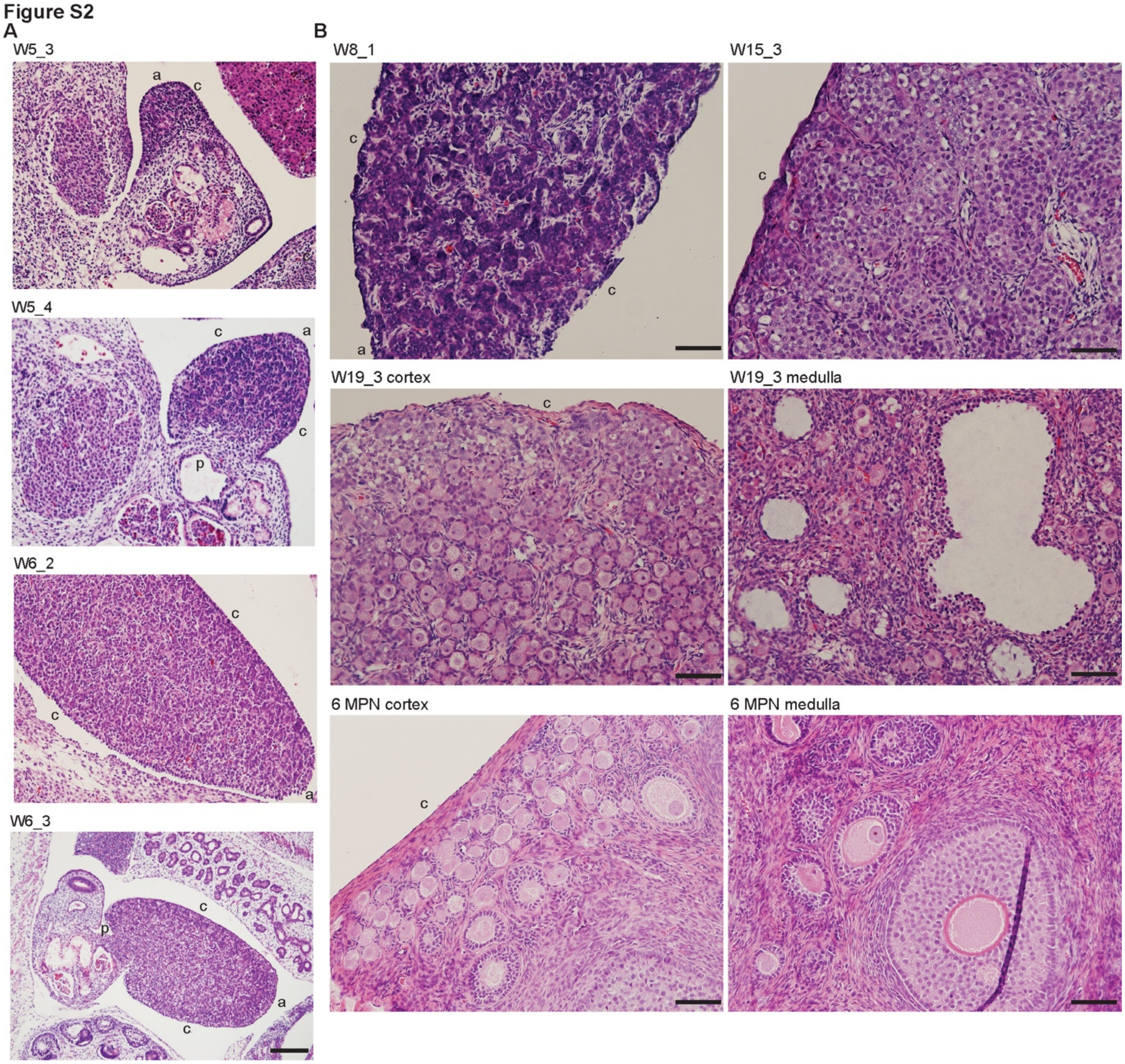

**Fig. S2. associated with Fig.1. (A)** Higher magnification H&E images of W5 and W6 gonads from torso section. Right gonad for W5_3, left gonad for W5_4 and W6_3; only one gonad is in plane for W6_2. **(B)** Higher magnification H&E images of W8, W15, W19 and 6 MPN gonads, with separate cortex and medulla images at W19 and 6 MPN. Labels indicate orientation – cortex (c), anterior (a), posterior (m). All scale bars 50 µM.

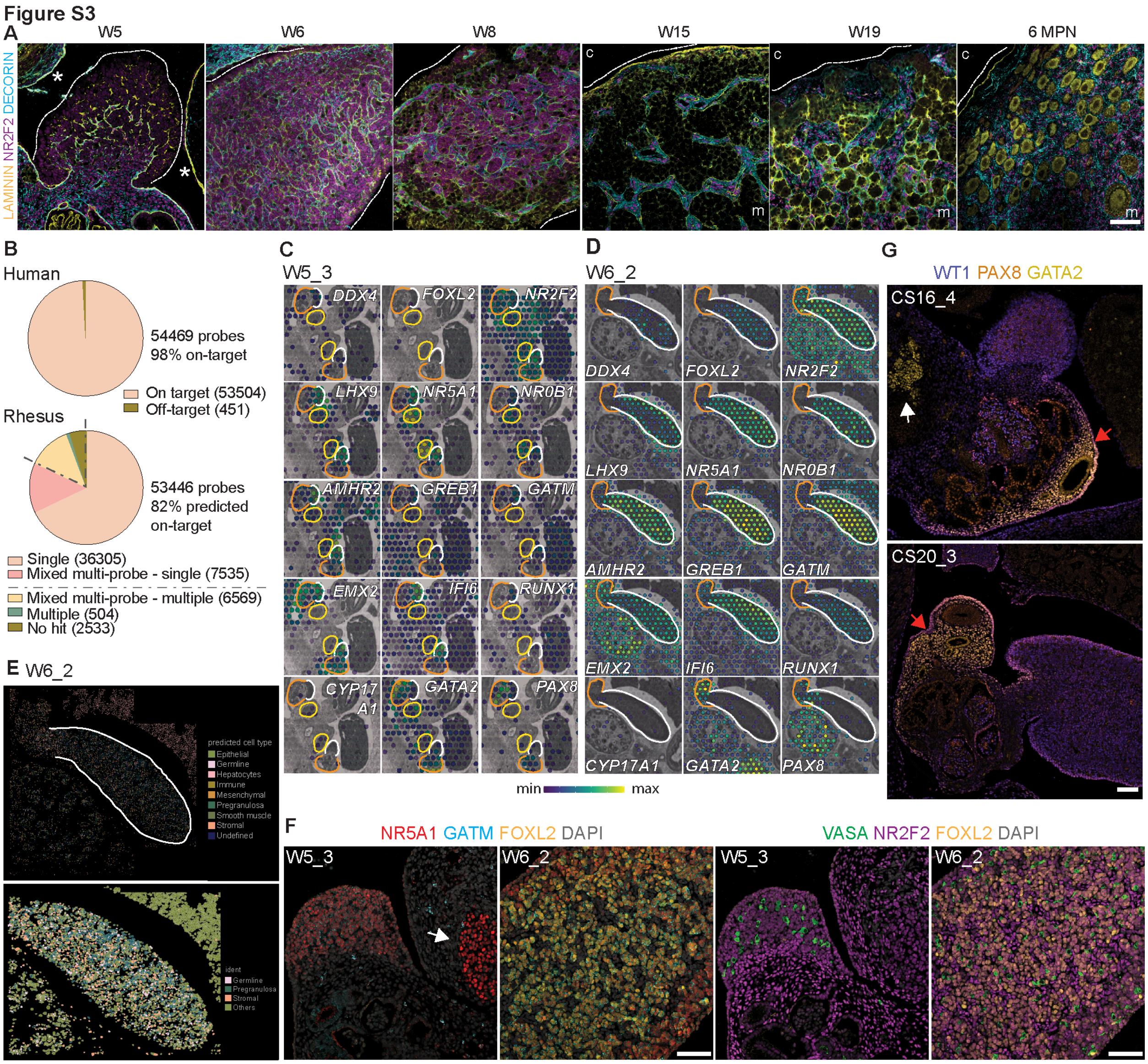

**Fig. S3. associated with Fig. 1 and 2.** Immunofluorescence analysis in W5, W6, W8, W15, W19 and 6 MPN rhesus ovaries for stromal (NR2F2, magenta) and extracellular matrix component markers (laminin, yellow; Decorin, cyan). Nuclei counterstained with DAPI (grey). Labels indicate orientation – cortex (c), medulla (m). White dashed line outlines outer cortex; stars highlight contiguous laminin deposition on adjacent tissue in contrast to ovary at W5. Scale bars 50 µM. **(B)** Pie chart illustrating *in silico* predicted coverage of the Visium Human Whole Transcriptome probe set for the human (left) and rhesus macaque (right) genomes. **(C)(D)** Spatial distribution of genes of interest mapped onto a gray scale image of the H&E stained section in W5_3 and W6_2 tissues used for Visium CytAssist spatial transcriptomics, cropped to focus on gonadal, mesonephric and adrenal regions. Ovary (white outline), mesonephros (orange) and adrenal (yellow) regions are outlined; scale colour from blue (min) to yellow (max) represents the expression level. **(E)** CosMx spatial molecular imager (SMI) analysis of W6_2 showing spatial distribution of all annotated cell clusters in the selected fields of view (top panel) and cell segmentation overlay showing distribution of germline, pregranulosa and stromal cells (bottom panel, additional clusters marked as “Others”). White dashed line indicates ovary region. **(F)** Immunofluorescence analysis for WT1 (blue), PAX8 (orange) and GATA2 (yellow) in W5_2 and W6_3 tissues. Arrows indicate GATA2 mesonephric (red arrows) or adrenal (white) expression. Scale bars 50 µM. **(G)** Immunofluorescence analysis for known genes (VASA, green; NR2F2, magenta; FOXL2, yellow) and genes identified in in the DEG analysis (NR5A1, red; GATM, cyan) in W5 and W6 ovaries. White arrow indicates NR5A1+ cells in the adrenal gland in W5_3 top panel. Scale bars 50 µM.

**Figure S4**

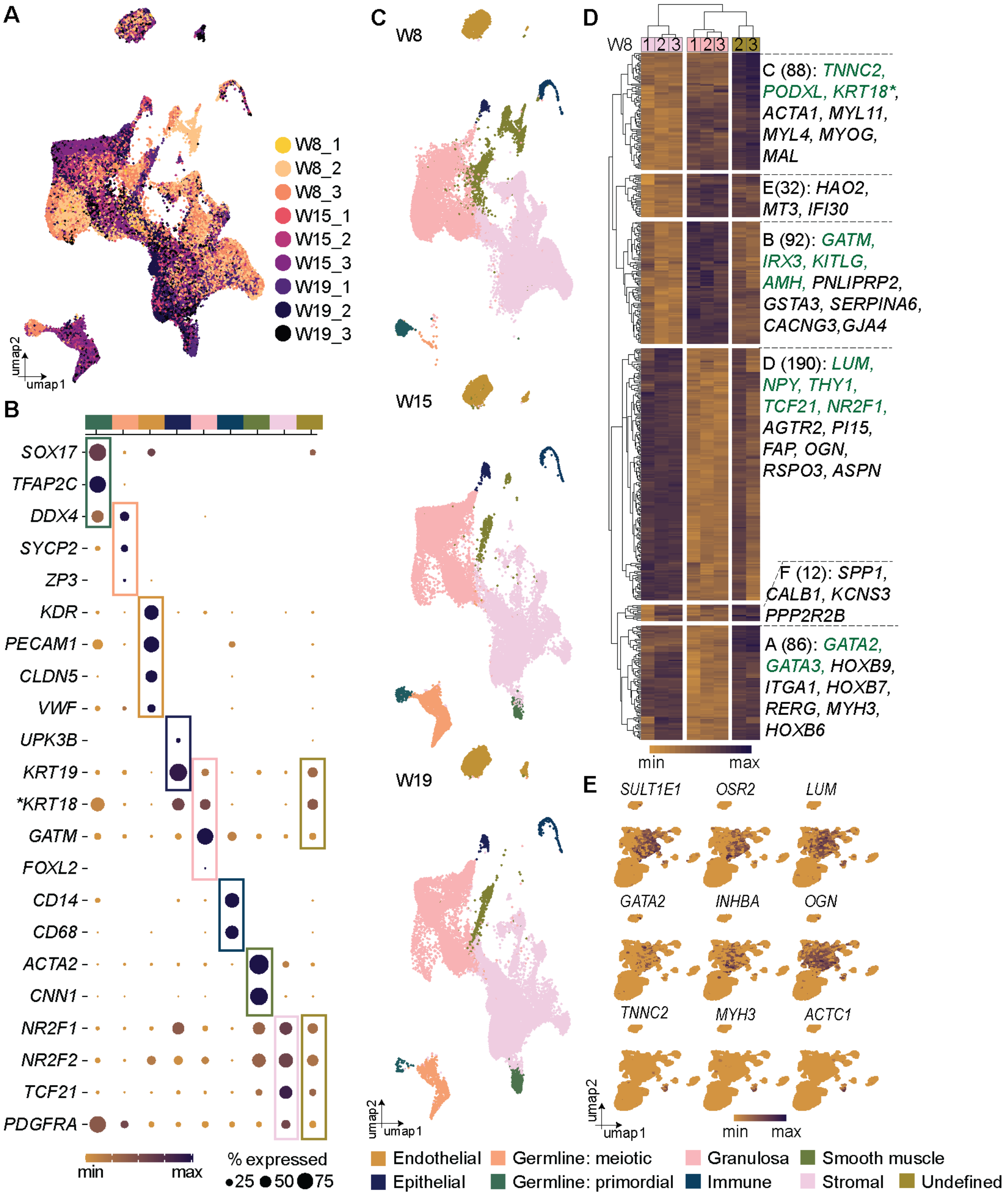

**Fig. S4. associated with Fig. 3. (A)** UMAP plot showing the distribution of single cells collected from rhesus foetal ovaries at W8, W15 and W19 colored based on their donor of origin (n=3 biological replicates per timepoint). **(B)** Dot plot comparing percentage expression of selected lineage-associated genes used to annotate Seurat clusters by cell type in Fig. 3A. Expression is plotted on a high-to-low scale (indigo-yellow) with dot size reflecting the percentage of cells within a cluster that express a given gene. **(C)** UMAP plot from Fig. 3A split according to timepoint. Granulosa, stromal and undefined cell sub-clusters have been collapsed into single colors. **(D)** Heatmap of top 500 most variable genes following pseudobulk pairwise comparisons of granulosa, stromal and undefined cell clusters at W8 (K-means clustering). Columns/rows are arranged according to hierarchical clustering; each row represents a gene, and each column represents a single donor. Gene expression is plotted according to log 2 normalized counts per million on a high-to-low scale (indigo-yellow). Stromal (purple), granulosa (pink) and undefined (green) column key is included at the top. Selected genes from each cluster are listed on the right of the heatmap; known granulosa or stromal associated genes in green and topmost enriched identified genes in black. **KRT18 = ENSMMUG00000031911*. See also Data S2. **(E)** Expression of additional markers enriched in the ovary from the DEG analysis at W5 and W6 cast on the UMAP plot from Fig. 3C. Normalized expression is plotted on a high-to-low scale (indigo-yellow).

Figure S5

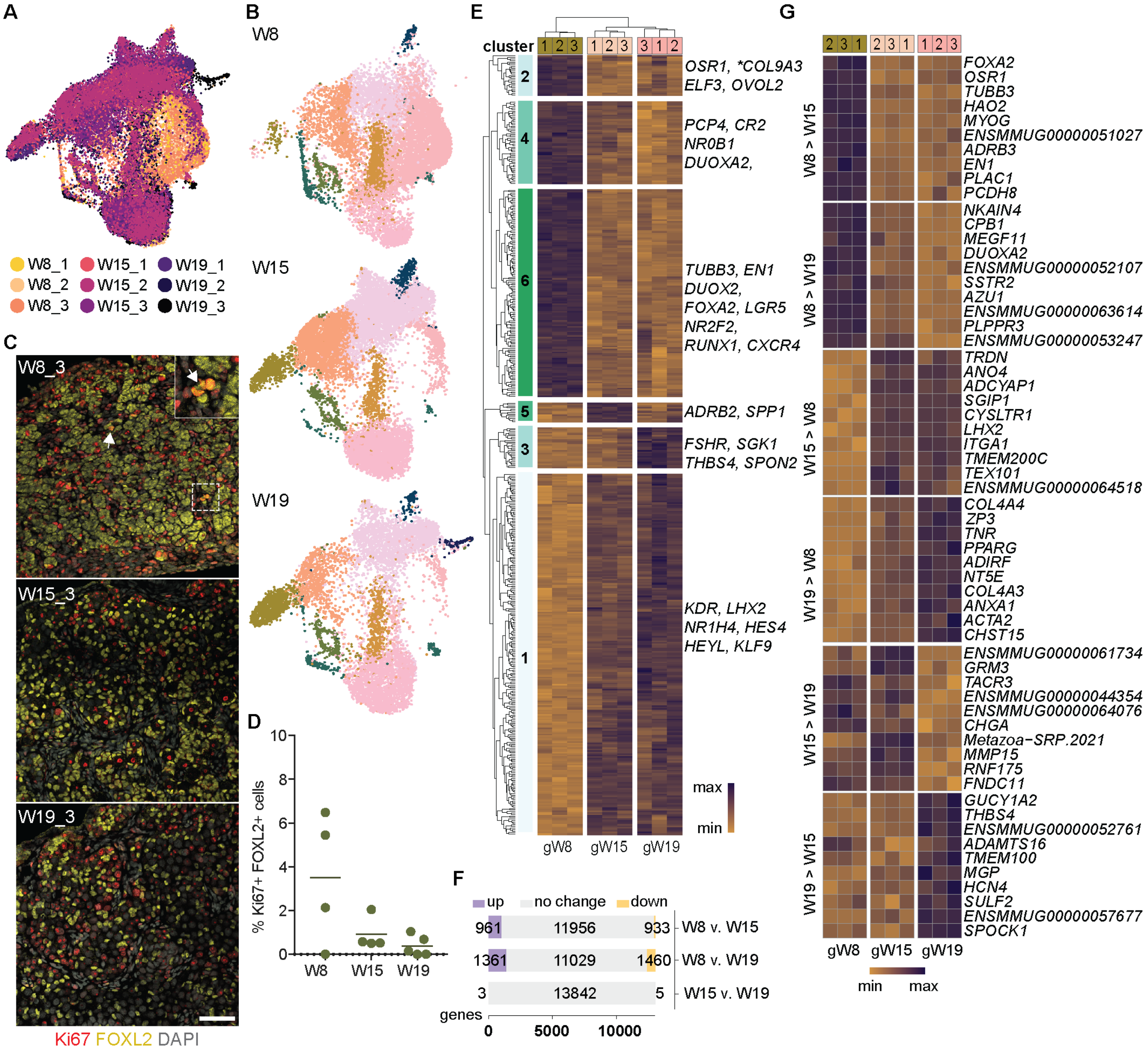

**Fig. S5 associated with Fig. 3. (A)** UMAP plot showing the distribution of single cells in the granulosa subset coloured based on their donor of origin (n=3 biological replicates per timepoint). **(B)** UMAP plot from Figure 3E split according to timepoint. **(C)** Immunofluorescence analysis for Ki67 (red) and FOXL2 (yellow) in W8, W15 and W19 ovaries. Arrows indicate overlapping expression; magnified area (dashed) shown in square inset. Scale bars 50 µM. (**D**) Quantification of the percentage of FOXL2+ cells at W8, W15 and W19 that express Ki67 by immunofluorescence. **(E)** Heatmap of top 500 most variable genes following pseudobulk pairwise comparisons of granulosa cells at W8, W15 and W19 (K-means clustering Columns/rows are arranged according to hierarchical clustering; each row represents a gene, and each column represents a single donor. Gene expression is plotted according to log 2 normalized counts per million on a high-to-low scale (indigo-yellow). W8 (green), W15 (peach) and W19 (pink) column key is included at the top. **(F)** Bar graph showing number of upregulated, downregulated or unchanged genes following pseudobulk pairwise comparisons. See also Data

S3. **(G)** Heatmap of top 10 enriched genes (FDR adjust p-value >0.05, log2 fold change >1 or <-1) in granulosa cells following pseudobulk differential gene expression analysis; corresponding pairwise comparison is noted on the left y-axis. Gene expression is plotted on a high-to-low scale (indigo-yellow). Stromal (purple), granulosa (pink) and undefined (green) column key is included at the top. Selected topmost enriched genes from each cluster are listed on the right of the heatmap

Figure S6

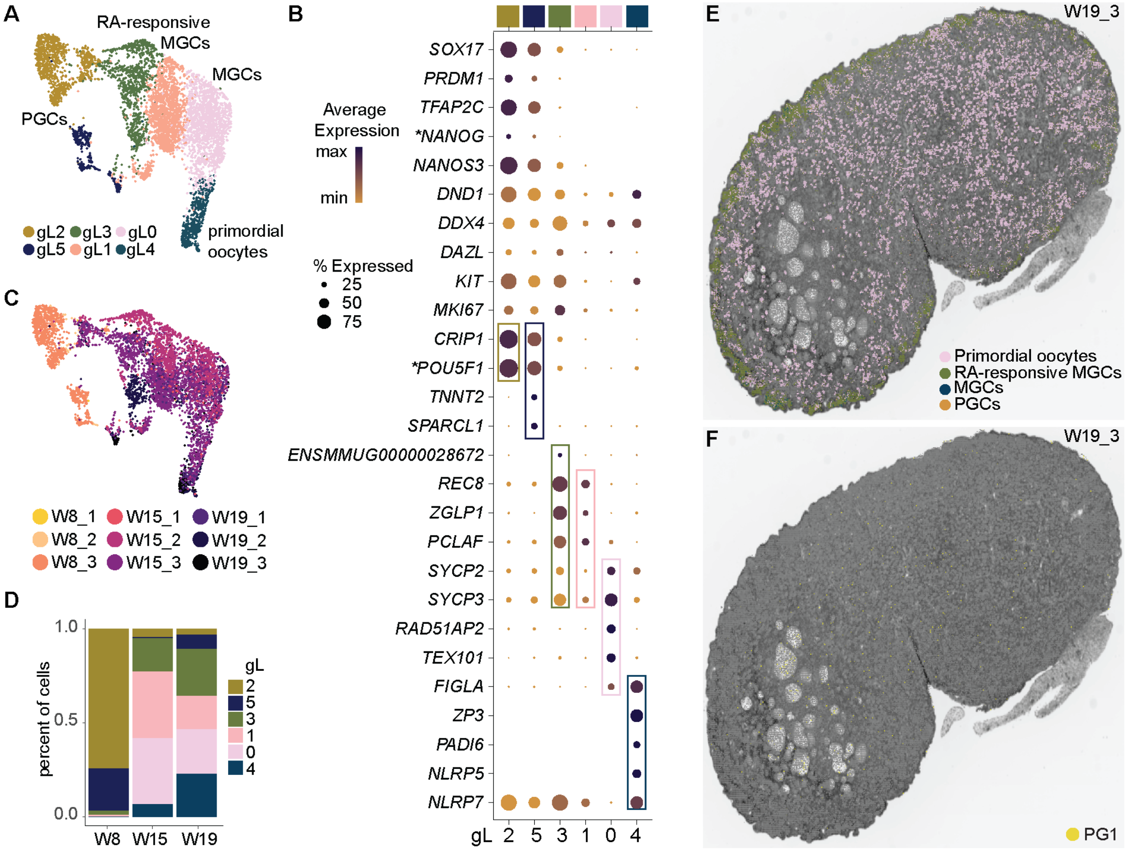

**Figure S6, associated with Figure 4. (A)** UMAP plot of germline subset (clusters a6 and a12 from Fig.3A) re-clustered and colored according to Seurat clusters**. (B)** Dot plot of known germline-associated genes, or genes identified as highly enriched in each cluster (rectangles). Expression plotted on a high-to-low scale (indigo-yellow); dot size reflects percentage of cells expressing given gene. **NANOG = ENSMMUG00000032158*, **POU5F1 = ENSMMUG00000015688*. **(C)** UMAP plot showing the distribution of single cells in the germline subset colored based on their donor of origin (n=3 biological replicates per timepoint). **(D)** Bar graph of germline sub-cluster proportions at each time point. **(E,F)** Spatial distribution of germline bins (E) and granulosa PG1 subcluster bins identified in Visium CytAssist HD analysis of W19_3 mapped onto a black and white H&E image. See Methods for details.

Figure S7

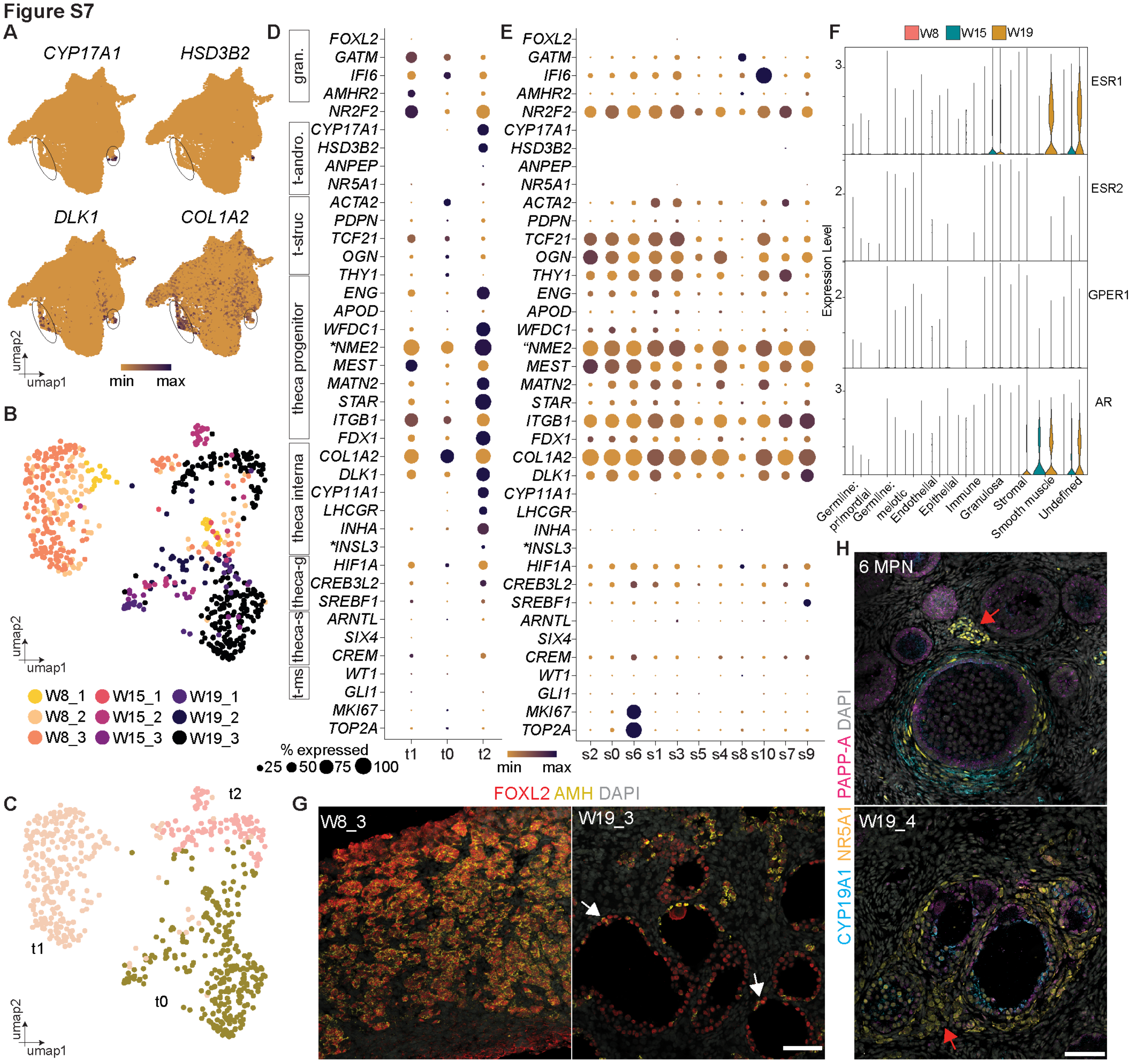

**Figure S7, associated with Figure 5. (A)** Expression of theca-associated markers enriched in cluster g7 (ovals) cast on the UMAP plot from Fig. 3E. Normalized expression is plotted on a high-to-low scale (indigo-yellow). **(B)** UMAP plot showing the distribution of single cells in the re-clustered g7 subset colored based on their donor of origin (n=3 biological replicates per timepoint). **(C)** UMAP plot from Fig. S7B colored based on Seurat analysis designated clusters. **(D)** Dot plot comparing percentage expression of selected granulosa-associated (*FOXL2, GATM*), stromal (*NR2F2*) and theca-associated genes identified in the literature in each of the Seurat clusters indicated in Fig. S7C. Genes for androgenic (t-andro) and structural theca (t-struc)(*75*); theca progenitor(*73, 74*); theca interna(*74, 76*); Theca-G and Theca-S(*78*); mouse theca (t-ms)(*77*). Expression is plotted on a high-to-low scale (indigo-yellow) with dot size reflecting the percentage of cells within a cluster that express a given gene. **NME2 = ENSMMUG00000001940*; **INLS3 = ENSMMUG00000042375*. **(E)** Dot plot comparing percentage expression of theca-associated genes in stromal clusters from the wider dataset (clusters a0, a2, a10 and a5 from Figure 3A extracted and re-clustered). **(F)** Violin plot of expression of estrogen receptors (*ESR1, ESR2, GPER*) and androgen receptor (*AR*) in the 10x Chromium dataset clusters, split according to timepoint. **(G):** Immunofluorescence analysis for AMH (yellow) and FOXL2 (red) in W8 and W19 ovaries. Scale bars 50 µM. **(H)** Immunofluorescence analysis for CYP17A1 (cyan), NR5A1 (yellow) and PAPP-A (magenta) in 6 MPN and W19 ovaries. Scale bars 50 µM.

Figure S8

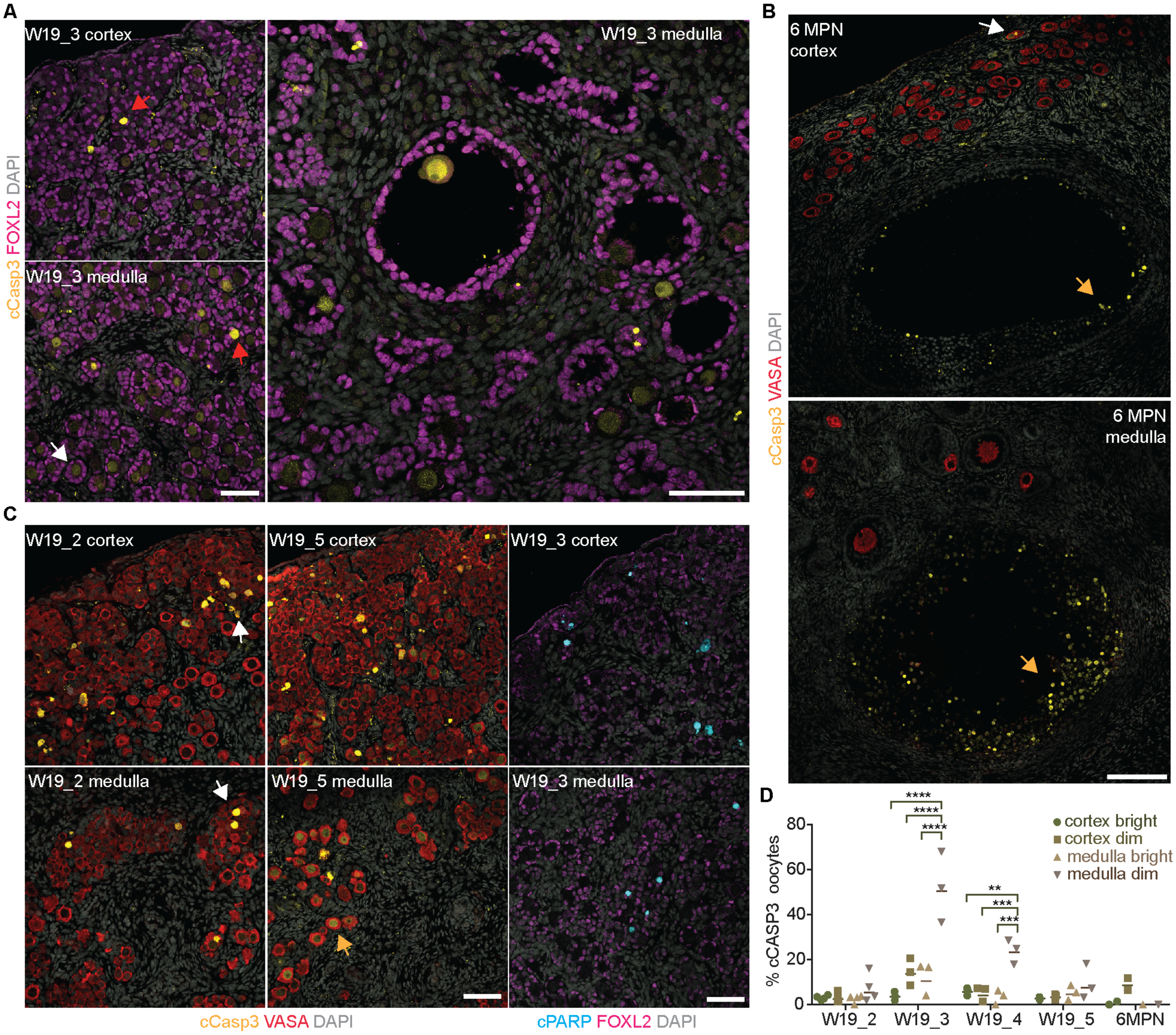

**Figure S8, associated with Figure 5. (A)** Immunofluorescence analysis for FOXL2 (magenta), cleaved Caspase-3 (yellow) expression and DAPI (grey) at W19. **(B,C)** Immunofluorescence analysis for VASA (red), cleaved Caspase-3 (yellow) expression and DAPI (grey) at W19 and 6 MPN, and cleaved PARP (cyan) and FOXL2 (magenta) at W19. All scale bars 50 μM. **(D)** Quantification of the percentage of oocytes at W19 or 6MPN that express cleaved Caspase3 by immunofluorescence. ****p=0.0001, ***p=0.001, **p=0.01 (two-way ANOVA).

**Table S1. Gonad sample key.**

| Manuscript ID | Animal ID / Library Name | Genotype | Library ID (Technical Replicates=A, B) | 10x Chromium | 10x Visium | 10x Visium HD | NanoString CosMx | IF images |
|---|---|---|---|---|---|---|---|---|
| W5_1 | ONPRC022 | XY (Sox9+) | ONPRC022 | - | X | - | - | - |
| W5_2 | ONPRC029 | XY (Sox9-) | ONPRC029 | - | X | - | - | - |
| W5_3 | ONPRC031 | XX (Foxl2-) | ONPRC031 | - | X | - | - | X |
| W5_4 | ONPRC032 | XX (Foxl2+) | ONPRC032 | - | X | - | - | X |
| W6_1 | ONPRC019 | XY | ONPRC019 | - | X | - | - | - |
| W6_2 | ONPRC020 | XX | ONPRC020 | - | X | - | X | X |
| W6_3 | ONPRC021 | XX | ONPRC021 | - | X | - | X | X |
| W8_1 | ONPRC018 | XX | ONPRC018_A<br>ONPRC018_B | X<br>X | - | - | - | X |
| W8_2 | ONPRC025 | XX | ONPRC025_A<br>ONPRC025_B | X<br>X | - | - | - | - |
| W8_3 | ONPRC027 | XX | ONPRC027_A<br>ONPRC027_B | X<br>X | - | - | - | - |
| W15_1 | ONPRC006 | XX | ONPRC006_A<br>ONPRC006_B | X<br>X | - | - | - | - |
| W15_2 | ONPRC007 | XX | ONPRC007_A<br>ONPRC007_B | X<br>X | - | - | - | - |
| W15_3 | ONPRC012 | XX | ONPRC012_A<br>ONPRC012_B | X<br>X | - | - | - | X |
| W19_1 | ONPRC001 | XX | ONPRC001_A<br>ONPRC001_B | X<br>X | - | - | - | - |
| W19_2 | ONPRC016 | XX | ONPRC016_A<br>ONPRC016_B | X<br>X | - | - | - | X |
| W19_3 | ONPRC017 | XX | ONPRC017_A<br>ONPRC017_B | X<br>X | - | X | - | X |
| W19_4 | 29906 | XX | - | - | - | - | - | X |
| W19_5 | 30466 | XX | - | - | - | - | - | X |
| 6 MPN | 26861 | XX | 26861 | - | - | - | - | X |

ONPRC017_A library excluded from downstream analysis as did not pass QC metrics.

### Table S2. RNA sequencing analysis statistics.

| 10x Chromium | | | | | | | | |
|---|---|---|---|---|---|---|---|---|
| Manuscript ID | Library ID | Raw reads | %Mapped | Estimated cell no. | Median genes / cell | UMI / cell | Valid cells | Totals |
| **W8_1** | ONPRC018_A | 310,770,130 | 86.5 | 908 | 2,245 | 5,796 | 757 | |
| | ONPRC018_B | 277,228,442 | 86.3 | 747 | 1,848 | 4,611 | 664 | |
| **W8_2** | ONPRC025_A | 283,601,942 | 89.1 | 9,832 | 1,964 | 4,492 | 7,675 | |
| | ONPRC025_B | 330,417,080 | 89.0 | 13,796 | 1,801 | 3,967 | 10,626 | |
| **W8_3** | ONPRC027_A | 283,073,916 | 90.0 | 10,740 | 1,850 | 3,982 | 8,428 | |
| | ONPRC027_B | 269,567,924 | 89.4 | 12,348 | 1,713 | 3,570 | 9,545 | 37,695 |
| **W15_1** | ONPRC006_A | 133,476,515 | 87.2 | 1,669 | 2,482 | 6,590 | 1,275 | |
| | ONPRC006_B | 199,122,808 | 87.6 | 1,767 | 2,566 | 6,945 | 1,380 | |
| **W15_2** | ONPRC007_A | 109,388,009 | 88.5 | 6,002 | 1,522 | 2,968 | 5,153 | |
| | ONPRC007_B | 96,095,328 | 88.1 | 6,251 | 1,413 | 2,740 | 5,305 | |
| **W15_3** | ONPRC012_A | 290,181,733 | 73.2 | 8,812 | 1,489 | 3,204 | 7,028 | |
| | ONPRC012_B | 278,107,923 | 83.5 | 14,457 | 1,334 | 2,919 | 10,573 | 30,714 |
| **W19_1** | ONPRC001_A | 238,602,610 | 87.8 | 5,036 | 1,544 | 3,546 | 3,817 | |
| | ONPRC001_B | 255,106,102 | 88.1 | 5,592 | 1,520 | 3,482 | 4,104 | |
| **W19_2** | ONPRC016_A | 240,956,237 | 86.2 | 7,977 | 1,777 | 4,060 | 6,438 | |
| | ONPRC016_B | 260,764,653 | 86.1 | 8,265 | 2,054 | 4,955 | 6,501 | |
| **W19_3** | ONPRC017_A | 253,359,275 | 85.7 | n/a (excluded) | n/a (excluded) | n/a (excluded) | n/a (excluded) | |
| | ONPRC017_B | 284,320,096 | 84.9 | 13,252 | 1,393 | 2,894 | 9,978 | 30,838 |
| 99,247 cells in final Seurat object | | | | | | | | |

| 10x Visium | | | | | | | | |
|---|---|---|---|---|---|---|---|---|
| Manuscript ID | Library ID | Alternate ID | No. of spots | No. of reads | Mean reads / spot | Median genes / spot | % mapped to probe set | Genes detected |
| **W5_1** | ONPRC022 | emb_22 | 1,293 | 109,070,458 | 84,355 | 6,765 | 90.0 | 16,339 |
| **W5_2** | ONPRC029 | emb_29 | 984 | 100,240,074 | 101,870 | 7,890 | 88.1 | 16,356 |
| **W5_3** | ONPRC031 | emb_31 | 780 | 89,964,490 | 115,339 | 7,740 | 89.3 | 16,322 |
| **W5_4** | ONPRC032 | emb_32 | 1,079 | 94,739,013 | 87,803 | 7,243 | 87.6 | 16,391 |
| **W6_1** | ONPRC019 | emb_19 | 1,976 | 131,936,299 | 66,769 | 7,538 | 88.9 | 16,371 |
| **W6_2** | ONPRC020 | emb_20 | 2,824 | 162,974,686 | 57,711 | 7,048 | 88.8 | 16,393 |
| **W6_3** | ONPRC021 | emb_21 | 3,640 | 306,360,197 | 84,165 | 7,396 | 86.0 | 16,428 |

| 10x Visium HD | | | | | | | | |
|---|---|---|---|---|---|---|---|---|
| Manuscript ID | Library ID | Alternate ID | No. of 8 µm bins | No. of reads | Mean reads / bin | Median UMIs / bin | % mapped to probe set | Genes detected |
| **W19_3** | ONPRC017 | emb_17 | 179,693 | 103,336,379 | 575.1 | 174.4 | 90.6% | 16,327 |

| Nanostring CosMx | | | | |
|---|---|---|---|---|
| Manuscript ID | Library ID | Alternate ID | Fields of view (FOVs) | Cells detected |
| **W6_2** | ONPRC020 | emb_20 | 17 | 20,312 |
| **W6_3** | ONPRC021 | emb_21 | 13 | 27,344 |

**Table S3. Primary antibodies.**

| **Primary Antibody** | **Concentration** | **Source** | **Catalogue no. / Identifier** |
|---|---|---|---|
| **anti-AMH*** | 1:100 | ABD Serotec | MCA2246T<br>RRID:AB_2226471 |
| **anti-AP2γ (TFAP2C)** | 1:200 | Santa Cruz | sc-12762<br>RRID:AB_667770 |
| **anti-cleaved Caspase3** | 1:200 | Cell Signaling | 9661<br>RRID:AB_2341188 |
| **anti-CK18** | 1:200 | Abcam | ab7797<br>RRID:AB_306086 |
| **anti-CK19** | 1:200 | Novus Biologicals | NBP2-15186<br>RRID: AB_2737392 |
| **anti-cKIT** | 1:200 | DAKO | A4502 |
| **anti-cleaved PARP** | 1:200 | Cell Signaling | 5625<br>RRID:AB_10699459 |
| **anti-CYP17A1** | 1:200 | Santa Cruz | SC-374244<br>RRID:AB_10988393 |
| **anti-CYP19A1** | 1:200 | Novus Biologicals | NB100-1596<br>RRID:AB_10000919 |
| **anti-Decorin** | 1:200 | R&D Systems | AF143<br>RRID:AB_354790 |
| **anti-GATA2** | 1:100 | Abcam | ab109241<br>RRID: AB_10865130 |
| **anti-GATA4*** | 1:200 | Abcam | ab84593<br>RRID:AB_10670538 |
| **anti-GATM** | 1:200 | Abcam | ab119269<br>RRID:AB_10902241 |
| **anti-FOXL2** | 1:100 | Novus Biologicals | NB100-1277<br>RRID:AB_2106188 |
| **anti-Ki67 (mouse)** | 1:200 | BD Biosciences | 556003<br>RRID: AB_396287 |
| **anti-Ki67 (rat)** | 1:200 | Invitrogen | 14-5698-80<br>RRID:AB_10853185 |
| **anti-Laminin** | 1:100 | Abcam | ab11575<br>RRID: AB_298179 |
| **anti-NR2F2** | 1:200 | R&D Systems | PP-H7147-00<br>RRID:AB_2155627 |
| **anti-NR5A1** | 1:200 | R&D Systems | PP-N1665-00<br>RRID:AB_2251509 |
| **anti-PAPP-A** | 1:100 | R&D Systems | AF2487SP<br>RRID:AB_2159346 |
| **anti-PAX8** | 1:100 | Novus Biologicals | NBP2-29903 |
| **anti-PRDM1** | 1:100 | Cell Signaling | 9115S<br>RRID:AB_2169699 |
| **anti-SOX17** | 1:500 | R&D Systems | AF1924<br>RRID: AB_355060 |
| **anti-VASA (rabbit)** | 1:200 | Abcam | ab13840<br>RRID: AB_443012 |
| **anti-VASA (goat)** | 1:200 | R&D Systems | AF2030<br>RRID: AB_2277369 |
| **anti-WT1** | 1:100 | Abcam | ab89901<br>RRID:AB_2043201 |

*Antigen retrieval with sodium citrate buffer

### Table S4. Secondary antibodies.

| Secondary Antibody (all at 1:400) | Source | Catalogue no. / Identifier |
|---|---|---|
| donkey anti-goat IgG AF488 | Jackson Labs | 705-546-147<br>RRID:AB_2340430 |
| donkey anti-goat IgG AF647 | ThermoFisher | 705-605-147<br>RRID:AB_2340437 |
| goat anti-mouse $IgG_{2a}$ AF488 | ThermoFisher | 715-546-150<br>RRID: AB_2535771 |
| donkey anti-mouse IgG AF594 | Jackson Labs | 715-585-150<br>RRID:AB_2340854 |
| donkey anti-rabbit IgG AF488 | Jackson Labs | 711-545-152<br>RRID:AB_2313584 |
| donkey anti-rabbit IgG AF647 | Jackson Labs | 711-605-152<br>RRID:AB_2492288 |
| donkey anti-rat IgG AF647 | Jackson Labs | 712-606-153<br>RRID:AB_2340696 |
| donkey anti-goat IgG AF488 | Jackson Labs | 705-546-147<br>RRID:AB_2340430 |

**Data S1. (separate file)**
Tables associated with Fig.2 and Fig. S3.
A: Differential gene expression (DEG) analysis tables from Visium CytAssist analysis comparing W5 ovaries (W5_3, W5_4) to mesonephros and adrenal regions and to all tissues on the section. Threshold: pval_adj<0.05, avg-log2FC > 1.
B: CytAssist DEG comparisons of W6 ovaries (W6_2, W6_3) to mesonephros and adrenal regions and to all tissues on the section. Threshold: pval_adj <0.05, avg-log2FC > 1.
C: CytAssist DEG comparisons of W5 ovaries to each other and to W6 ovaries. Threshold: pval_adj <0.05, avg-log2FC > 1.
D: CytAssist DEG comparisons of W5 and W6 ovaries to W5 testes (W5_1, W5_2) and W6 testis (W6_1). Threshold: pval_adj <0.05, avg-log2FC > 1.
E: Cluster analysis of NanoString CosMx cell type clusters from W6 sections. Threshold: pval_adj <0.05.

**Data S2. (separate file)**
Tables associated with Fig.3, and Fig. S5.
A: DEG comparisons following pseudobulk analysis of W8 stroma, granulosa and undefined cell clusters. Threshold: FDR<0.05 in at least one comparison.
B: DEG comparisons following pseudobulk analysis of granulosa cells at W8, W15 and W19. Threshold: FDR<0.05 in at least one comparison.

**Data S3. (separate file)**
Tables associated with Fig.3, Fig. S5 and S7.
A: Genes in granulosa clusters from UMAP plot in Fig. 3E. Threshold: pval_adj <0.05.
B: Genes in theca clusters from UMAP plot in Fig. S7D. Threshold: pval_adj <0.05.
